# AB_SA: Tracing the source of bacterial strains based on accessory genes. Application to *Salmonella* Typhimurium environmental strains

**DOI:** 10.1101/814459

**Authors:** Laurent Guillier, Michèle Gourmelon, Solen Lozach, Sabrina Cadel-Six, Marie-Léone Vignaud, Nanna Munck, Tine Hald, Federica Palma

## Abstract

The partitioning of pathogenic strains isolated in environmental or human cases to their original source is challenging. The pathogens usually colonize multiple animal hosts, including livestock, which contaminate food-producing and environment (e.g. soil and water), posing additional public health burden and major challenges in the identification of the source. Genomic data opens new opportunities for the development of statistical models aiming to infer the likely source of pathogen contamination. Here, we propose a computationally fast and efficient multinomial logistic regression (MLR) source attribution classifier to predict the animal source of bacterial isolates based on “source-enriched” loci extracted from the accessory-genome profiles of a pangenomic dataset. Depending on the accuracy of the model’s self-attribution step, the modeler selects the number of candidate accessory genes that better fit the model for calculating the likelihood of (source) category membership. The accessory genes-based source attribution (AB_SA) method was applied on a dataset of strains of *Salmonella* Typhimurium and its monophasic variants (*S*. 1,4,[5],12:i:-). The model was trained on 69 strains with known animal source categories (i.e., poultry, ruminant, and pig). The AB_SA method helped to identify eight genes as predictors among the 2,802 accessory genes. The self-attribution accuracy was 80%. The AB_SA model was then able to classify 25 over 29 *S.* Typhimurium and *S*. 1,4,[5],12:i:-isolates collected from the environment (considered as unknown source) into a specific category (i.e., animal source), with more than 85% of probability. The AB_SA method herein described provides a user-friendly and valuable tool to perform source attribution studies in few steps. AB_SA is written in R and freely available at https://github.com/lguillier/AB_SA.

**Author Notes:** All supporting data, code, and protocols have been provided within the article and through supplementary data files.

Supplementary material is available with the online version of this article.

**Abbreviations:** AB_SA, accessory-based source attribution; MLR, multinomial logistic regression; SNPs, single nucleotide polymorphisms; GFF, general feature format; AIC, Akaike information criteria.

**Data Summary:**

1. The AB_SA model is written in R, open-source and freely available Github under the GNU GPLv3 licence (https://github.com/lguillier/AB_SA).
2. All sequencing reads used to generate the assemblies analyzed in this study have been deposited in the European Nucleotide Archive (ENA) (http://www.ebi.ac.uk/ena) under project number PRJEB16326. Genome metadata and ENA run accession ID for all the assemblies are reported in the supplementary material.

**Impact Statement:** This article describes AB_SA (“Accessory-Based Source Attribution method”), a novel approach for source attribution based on “source enriched” accessory genomics data and unsupervised multinomial logistic regression. We demonstrate that the AB_SA method enables the animal source prediction of large-scale datasets of bacterial populations through rapid and easy identification of source predictors from the non-core genomic regions. Herein, AB_SA correctly self-attribute the animal source of a set of *S.* Typhimurium and *S*. 1,4,[5],12:i:- isolates and further classifies the 84% of strains contaminating natural environments in the pig category (with high probability ranging between ∼85 and ∼99%).

## Introduction

Tracing the origin of pathogenic microbial strains associated with human diseases or contamination of environmental settings is crucial for identifying targets for interventions in the food production chain from farm-to-fork. The process of estimating the probability that human cases or environmental contamination cases arise from putative sources of infection (i.e. animal reservoir, food product, and environmental) can be referred to as source attribution. A variety of methodological approaches has been developed for source attribution of foodborne pathogens: epidemiological approaches (e.g., outbreak data analysis, case-control/cohort studies), microbial subtyping methods, comparative exposure assessment, intervention studies, and expert elicitations (1; 2).

Some of the source attribution methods based on microbial sub-typing specifically consider genotypic data (3; 1). Genetic variations in microorganisms are the result of different evolutionary forces. These can be prompted by either neutral processes (genetic drift) or adaptive processes, such as the emergence of a competitively advantageous mutation in a given environment. Most bacterial populations are structured, i.e., their entirety does not form a genetically homogeneous unit, but rather consists of several distinct lineages or sub-lineages that are totally or partially isolated from one another. Factors such as geographical isolation, combined with random phenomena of genetic drift and sometimes with local adaptation, drives genetic differentiation.

When analysing genetic data for a given microorganism based on targets like a certain number and type of alleles, microsatellites, or single nucleotide polymorphisms (SNPs), the objective is often to detect whether these microorganisms are structured into different subpopulations, and if so, to identify the number of clusters, the strains composing them, and possible recombination events. Historical methods for studying the genetic relatedness of microbial populations are based on the reconstruction of phylogenetic trees from a matrix of proximities for each pair of strains, with these genetic proximities being typically calculated using the methods proposed by (4) or (5). Once the tree is built, it can be ‘cut’ at a certain point (e.g., after three levels of nodes from the root) to define the different clusters of strains (more or less equivalent to sub-types). Visually exploring the composition of the clusters (i.e., isolates from different backgrounds) provides a general overview for inferring sources and transmission. However, this approach has been rarely applied in source attribution as inference by phylogeny relies upon the robustness of the tree built on the genetic diversity between isolates, and strains to attribute and strains from sources are usually phylogenetically intermixed (6). Indeed, closely related strains can be found in multiple hosts challenging the association to a specific source by phylogenetic clustering (7). A particular case showing the utility of phylogenetic methods in the attribution of human salmonellosis to specific sources (e.g., turkey), by using epidemiological and genomic data, has been reported through the investigation of *Salmonella* Derby genetic diversity (8).

A much different approach relies on the assumption that genetic data (e.g., frequency of different allele numbers at a locus) can be explained by a probabilistic model whose parameters are unknown. Comparing genetic data (frequencies) among different strain populations make it possible to establish a link between them, e.g., between strains from human cases and different sources. Two structured population genetics models that are currently widely used for source attribution of foodborne diseases are the so-called STRUCTURE approach (9) and the Asymmetric Island Model (AIM) (10). These two models are based on different principles of genetic structuring of microbial populations, but the overall attribution approach is similar. These approaches have been successfully applied for source attribution of human sporadic strains for *Campylobacter* spp. (11; 12), *Salmonella* spp. and *Listeria monocytogenes (13)*.

Machine learning approaches are gaining interest in identifying the underlying genetic features associated with traits of microbial pathogens (14) and their use is also discussed for tracing the origin of an outbreak (15). For source attribution, recent studies consider this approach for predicting the source of sporadic human cases (16; 17; 18). In particular, (17) applied a Random Forest classifier for genomic source prediction. The authors revealed that 50 key genetic features were sufficient for robust source prediction of strains from *Salmonella* Typhimurium serovar. Most of the genetic features were accessory genes.

Among other supervised classification techniques, multinomial logistic regression (MLR) is a multi-class classification procedure. It is an extension of binary logistic regression allowing for more than two outcomes events. MLR has proven to provide a pertinent framework to carry out association analyses across multiple phenotypic traits (19; 20) and in foodborne outbreak investigations as a rule-out tool (21). Complex phenotypes, such as host adaptation in specific niches, have been linked to the presence of genes and genetic elements in some strains but not in others (referred as the “accessory genome”), mainly driven by horizontal DNA transfer (22; 7; 23). Association analysis on a pangenome scale may relate patterns of genotypic variation (e.g., the differential composition of accessory genes of multi-host lineages) to specific zoonotic niches. The objective of this study was therefore to study the performance of multinomial logistic regression in source attribution. In particular, the method was used for predicting to which animal reservoir will environmental strains of *Salmonella* Typhimurium (*S.* Typhimurium) and its monophasic variants (*S*. 1,4,[5],12:i:-) be attributed, given the variable set of source-enriched genes from the strains’ accessory genome.

## Materials and Methods

### Preliminary genomic analysis

High-quality assemblies of 98 bacterial isolates (see Äpplication to *Salmonella* Typhimurium and its monophasic variant genomes dataset”below) were generated by DTU FoodQCPipeline (https://bitbucket.org/RolfKaas/foodqcpipeline). In short, the FoodQCPipeline trimmed the raw reads using bbduk2. Reads were then *de novo* assembled using SPAdes v3.11.05 (24) in the last step of the pipeline. FastQC v0.11.5 (https://www.bioinformatics.babraham.ac.uk/projects/fastqc/) was applied in multiple steps of reads processing (e.g., before and after trimming), generating a quality control report for each sample. The quality of the *de novo* assemblies was finally assessed using Quast v4.5 (25). Besides, the maximum-likelihood phylogenetic reconstruction of the 98 genomes dataset was built on single-nucleotide polymorphisms (SNPs) identified in the core genome alignments to assess the applicability of the dataset in (26). The annotated tree shows that environmental isolates (i.e. unknown source) were intermixed with potential sources and the dataset is therefore eligible for source attribution.

### Accessory Based Source attribution (AB_SA) method

The Accessory Based Source attribution (AB_SA) method is based on genomics data. The method is a two-step process. First, the accessory genes enriched in the different sources are calculated (see “Preparation of input data for multinomial logistic model pangenome analysis” below). Then, a multinomial logistic regression is developed to predict the probability of animal sources membership on environmental isolates based on “source-enriched” accessory genes.

Multinomial regression is thus used to explain the relationship between one nominal dependent variable (with more than two levels), that is the source, and one or more independent variables, i.e. the enriched genes. For a source attribution situation with K sources, the multinomial regression model estimates *k* (*k* = *K -* 1) equations:

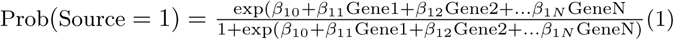

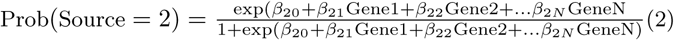

…

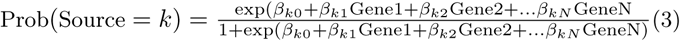

For the final source, the probability of association is derived from the K-1 other equations:

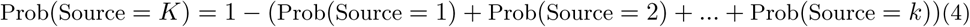

### Preparation of input data for multinomial logistic model pangenome analysis

The genomes assemblies were annotated in GFF3 format by using Prokka (Seemann, 2014). Roary (27) was used to determine the pangenome of the whole dataset of strains from the annotated assemblies. The dataset included both strains with known (i.e., animal reservoir) and unknown sources (i.e., environmental strains to attribute). To know which genes were enriched in each host groups, Scoary was used (28). Scoary takes as input the gene presence absence.csv file from Roary and a traits file reporting the source associated with each strain. Notice that the traits file is restricted to strains with known sources. The `--no_pairwise` and `--collapse` options from Scoary were applied to determine the genes that are enriched in each source. The `--no_pairwise` flag is used for enrichment analysis to avoid pairwise comparisons and perform population structure-naive calculation. The `--collapse` flag was used to merge genes that present the same pattern of distribution in the sources (a single gene, the first of each merged group of genes is then taken into account by the AB_SA method). Besides, the naive p-value is used to show the genes most overrepresented in a specific source. A naive threshold p-value of 0.01 was used to establish the list of potential genes of interest for the attribution of the source, sorted by strength of association per trait. To reduce or increase the number of genes to be considered by the AB_SA method, the user can modify the threshold p-value.

The `CreateInputMNL` function from the AB_SA method creates the input files for multinomial logistic model. It takes as arguments: the trait file used as input of Scoary and the gene presence/absence .Rtab from Roary, and the number of enriched genes to be taken in each source. The function returns two files: a file for fitting multinomial logistic model on data originating from source, and a file used for determining the probability of association the source of unknown strains (sporadic human strains or environmental strains).

### Training and testing of the multinomial logistic model

Association of sources and genes are then carried out by the multinomial model built on a combination of functions from the “caret”, “e1071” and “ROCR” R packages (29).

A split-sample approach was applied for training and testing the model using the dataset with known sources to select model tuning values and estimate the model performance through resampling. The dataset with known sources was randomly partitioned into complementary subsets in a ratio of 70:30, meaning that 70% of data will be used for model training and the remaining 30% for model testing (evaluating model performance) by (K-fold and bootstrapping) resampling method (30). The resampling method creates modified datasets of samples from the training sets and a model is fit to each resampled dataset to predict the corresponding set of hold-out samples. The aggregation of the results of each hold-out sample set is then used to estimate the resampling performance for finally assessing the more appropriate combination of tuning parameters to consider for the final model refit on the entire dataset.

The training and testing are carried by `MNLTrainTest` function in AB_SA. It takes as inputs the ouput file from `CreateInputMNL` function., the partition percentage (70:30 as default value) as well as the number of bootstrap.

### Assessment of the model’s performance

Trained models were assessed through accuracy metrics to select the optimal model. To avoid overfitting, the assessment of fitted logistic multinomial models was carried out with regularization. The Akaike Information Criteria (AIC) was used as a statistical measure of fit to penalize the number of parameters (predictor variables) included in the multinomial logistic model, helping to identify too complex models that tend to memorize training data. The AB_SA `MNLTrainTest` function returns both the density plot showing the performance estimates of accuracy and the AIC value.

### Prediction of strains with unknown source

Based on the accuracy of the trained model and the AIC values, the appropriate number of genes to include in the multinomial logistic model is selected by the modeler. The full set of strains with known sources is fitted with `MNLfit` function for these genes and samples with unknown source are then predicted using the AB_SA `MNLpredict` function, which returns a data frame with probability values for each animal source.

### Application to *Salmonella* Typhimurium and its monophasic variant genomes dataset

The dataset used to explore the relationship between genes of the accessory genome and the animal host is constituted by 98 strains belonging to *S.* Typhimurium and its monophasic variant and collected between 2010-2015. The dataset is fully described elsewhere (26) and reported in the Supplementary Material. The dataset is constituted of strains from known sources and strains with an unknown animal source (i.e. strains to attribute). The set of isolates with a known source comprises strains isolated from pigs (n=49), poultry (layers, broilers, turkeys, and ducks) (n=14) and ruminants (cattle, sheep, and goat) (n=6). For the strains to attribute, 29 strains of *Salmonella* were isolated from the environment (e.g., fresh or brackish water and soil) from both ANSES and IFREMER with the collaboration of the University of Caen (France). The IFREMER’s environmental isolates originate from a research project (31). They were isolated from freshwater (n=12) in Brittany and brackish water in Normandy (n=3). ANSES environmental isolates were isolated from soils (n=3) and freshwater (n=10). One strain isolated in crustacean was also associated with this environmental dataset (26).

## Results

### Pangenome-wide enrichment analysis of the *Salmonella* dataset

In total, 98 *S.* Typhimurium and monophasic variant strains were used in this study as input for implementing a multinomial logistic regression model of source attribution. Over 98 strains, 29 were isolated from environmental (i.e., water, soil samples, and a crab isolate) while the remaining 69 from animal sources (i.e., pigs, poultry, and ruminants). The whole dataset of genomes was used to extract the pangenome while only the genomes from animal sources were used to score accessory genes as enriched in each animal source. Over 6,988 genes composing the pangenome, 40% (n=2,802) represented the accessory genome (present in <99% of strains) and were then considered for the enrichment of genes in the animal sources. With a naive p-value<0.006, 10 genes were retained as enriched in the selected animal sources (Fig. 1, available in the Supplementary Material). However, a cluster of four correlated genes was collapsed into a merged unit. Only the first gene of the merged unit together with nine additional “source-enriched” genes will be considered as predictors by the AB_SA model. Most of the candidates’ genes are specifically enriched in a specific animal source (i.e. ruminants (n=3), pigs (n=2) and poultry (n=2)), while the remaining (n=3) are enriched in multiple sources (e.g., pigs and poultry).

**Figure 1:**
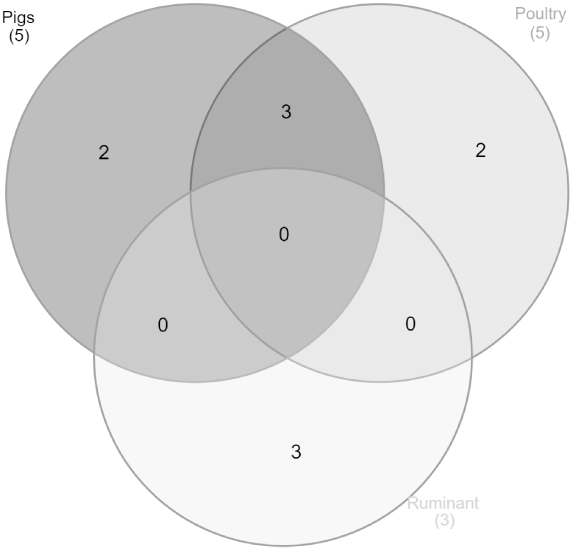
Number of enriched genes for the three animal sources

Feeding the MLR model with the source-enriched genes, a maximum number of genes to consider for predicting the source is arbitrarily selected. In order to select the optimal set of predictors, different numbers of genes (from 1 to 5) were tested, and for each case, accuracy and AIC were assessed (Table 1).

**Table 1:**
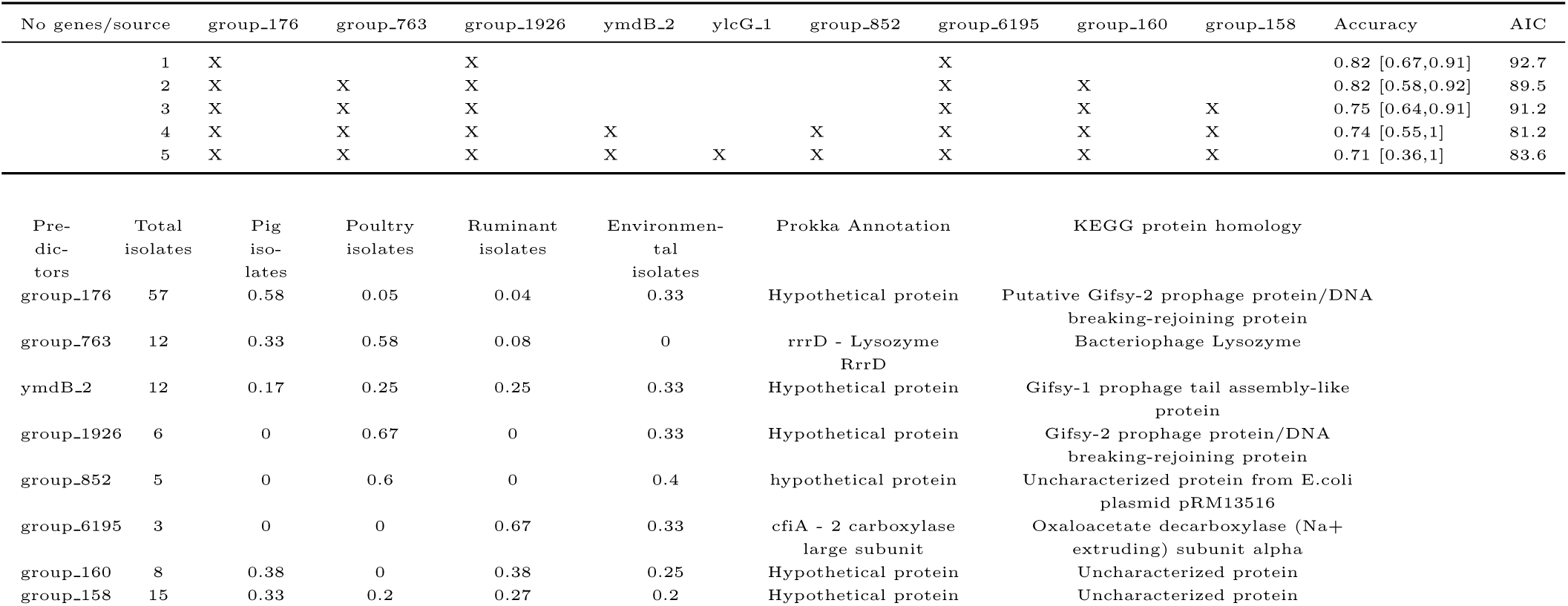
Tested multinomial logistic models with the accuracy obtained with the training and the AIC values

### Assessment of multinomial logistic model to predict the strains with known origins

For further performing an accurate animal source prediction, it is necessary to select the genes set that better discriminates pigs, poultry and ruminant-related genomes. When testing the ability to classify strains with known source by randomly selecting 70% and 30% of genomes for training and testing respectively, accuracy ranged from 0.71 to 0.82 according to the different genes included in the model (Table 1). Yet confidence intervals of the accuracies are large, and they could be considered as equivalent. AIC values help to distinguish the best model among the tested ones. In this study, the model including a total of eight genes as predictors (Table 1) was found to be the best model (with the lowest AIC value).

This set of “best” predictive genes is therefore used by the AB_SA model to classify genomes with unknown source. The relative importance of each predictor in estimating the model is calculated (Fig. 2). This statistical measure relates to the weight of each predictor in making a prediction, not whether or not the prediction is accurate. Fig. 2 presents the values of fitted *β*_*k*._ parameters. It shows that some genes have a higher weight than others. For example, group_6195 presence is strongly associated with bovine. In the same way, group_852 represents the highest coefficients for poultry.

**Figure 2:**
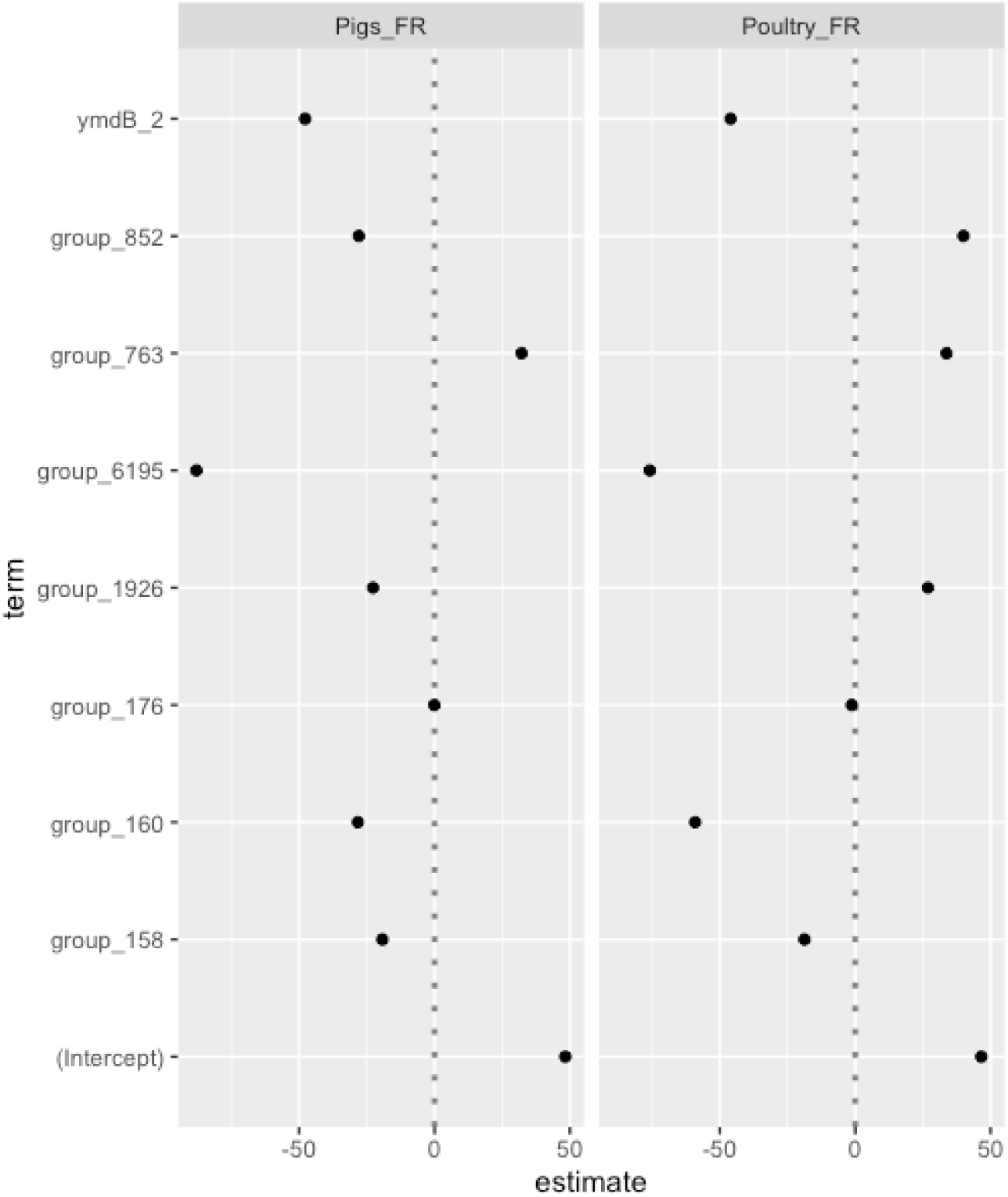
Estimates of the parameters of the multinomial logistic model. Ruminants _FR was the reference condition.

### Prediction of the origin of environmental strains

Fig. 3 shows the probabilities of the different environmental strains to be associated with one of the three sources. Six strains (i.e., 9, 12, 14, 25, 28, and 29) have a very high membership probability, that is, superior to 99%, to one of the three sources. The majority of the strains (n=19) has a high probability, ranging between 85-95%, of being associated to pig sources. For the four remaining strains (i.e., 5, 7, 11, and 24), the probability to be associated with a specific source is lower than 80% (e.g., ranging from ∼39% to ∼77%).

**Figure 3:**
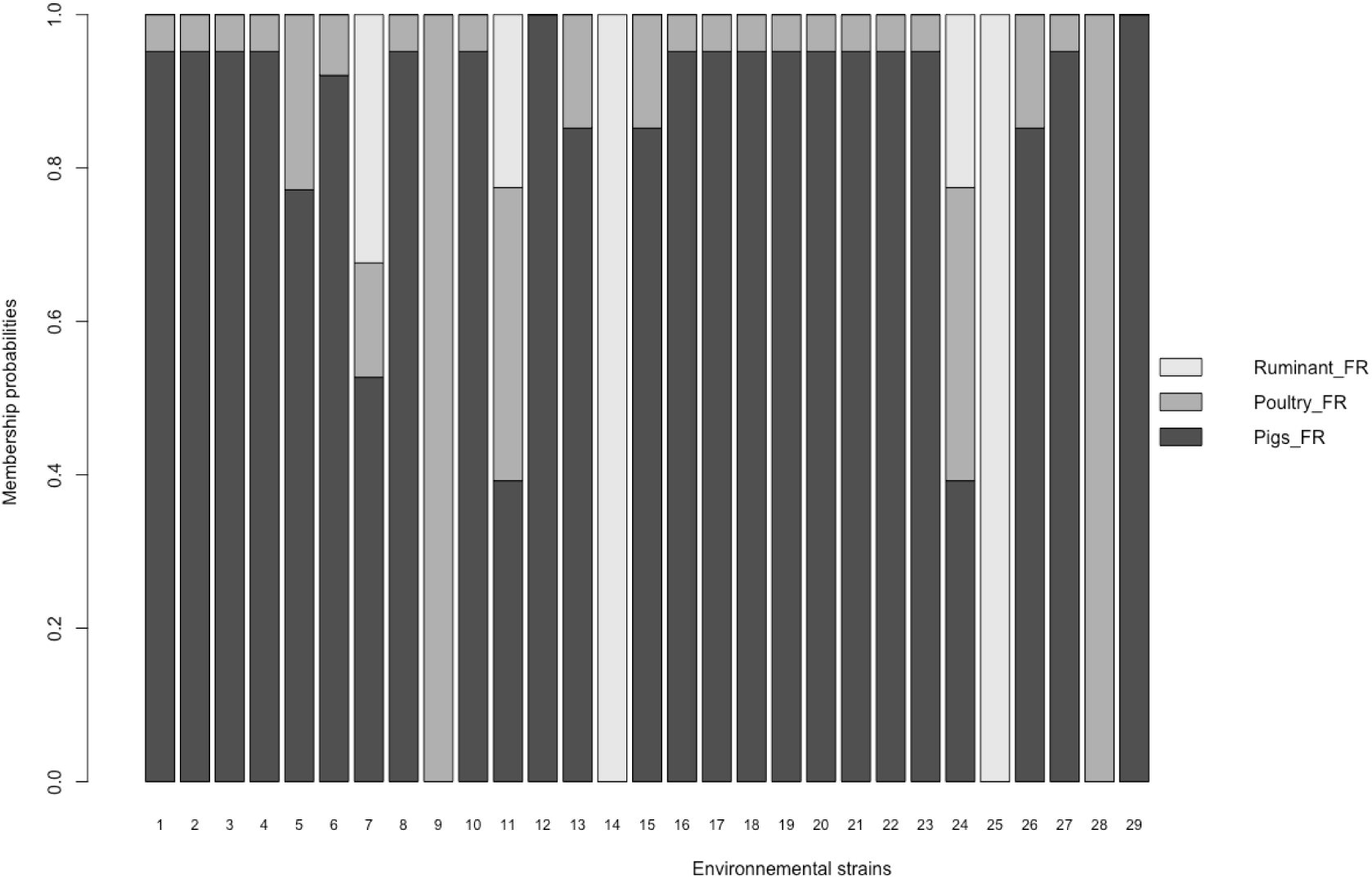
Individual source attribution of the 29 environmental strains of *Salmonella* Typhimurium to the three animal sources according to the ABSA method carried out on eight genes of the accessory genomes.

Fig. 4 shows the global attribution of the whole dataset of environmental strains to the three animal sources. For this dataset, pig source was found to be the main source of the strains isolated in the environment.

**Figure 4:**
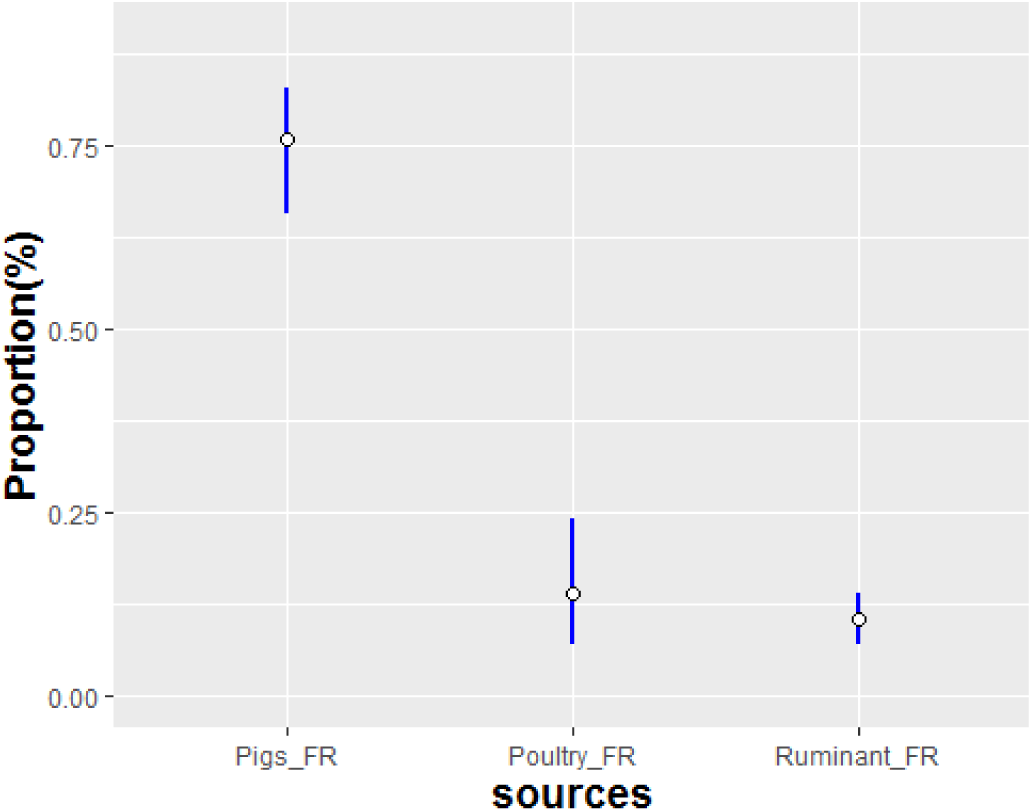
Global attribution of the environmental *Salmonella* Typhimurium strains to the three sources.

## Discussion

### Multinomial model for (WGS)-based source attribution

The environment is not a natural reservoir of *Salmonella* Typhimurium. This study thus focused on attribution of 29 *Salmonella* strains isolated from the environment (i.e. river and brackish water, soil, and crab) to potential sources using accessory based genes. The animal sources were grouped into three major categories (i.e., ruminants, poultry, and pigs) constituted by a dataset of 69 isolates of *Salmonella* Typhimurium and its monophasic variants. Genetic factors predictive of animal sources were identified in the accessory genome of isolates from this dataset through an innovative workflow based on enrichment and multinomial logistic analysis. After the selection of source-enriched genes as predictors, a maximum likelihood estimation is used by the AB_SA model to assess the probability of belonging to each animal source (categorical membership) for a given environmental isolate. The MLR-based models are less sensitive to data assumptions (e.g., normality, linearity and homogeneity of data) and have the advantage of limitation of overfitting, which is a common pitfall in machine learning approaches (32). It occurs when a complex model is trained on too few data points and becomes specific to the training data. As AB_SA returns both accuracy and AIC value (a penalty for model complexity), overfitting is prevented.

In addition, it is flexible in the choice of the level of significance (through threshold value for p-value) of enriched genes and in the number of candidate predictors per source to feed the multinomial logistic model. The MLR model also provides a measure of the weight of each predictor (i.e., gene) although the interpretation of the coefficients is not immediate. The interpretation might be further complicated by not having a single model, but as many models as the number of sources minus one.

Consistent with previously published source attribution studies, the overall self-attribution accuracy of this study was 75-80% (17; 18; 12).

### Number of genes predictive of animal source

The enrichment step of the AB_SA method allowed us to select the most relevant genes for modeling their effects with the multinomial logistic model. Within the 2,802 accessory genes, eight genes were finally selected in the best model for the prediction of the origin of environmental strains. The more predictors used in the training phase improved the model performance. However, using more than eight genes as predictors returns a worse AIC value in this dataset since the use of more data training leads to over-fitting in regression. This study confirms that high dimensional input data for source attribution models do not guarantee high performance. For example, (13) didn’t observe for *L. monocytogenes* an important improvement of accuracy of source attribution models when passing from 7-locus MLST to cgMLST.

### Source prediction of strains using accessory genes as predictors

In this study, eight genomic factors making up the ancillary genomes of the observed *Salmonella* dataset were selected as predictive of animal sources (Table 2, sequences available in the Supplementary Material) by the AB_SA method. Although the presence of several hypothetical proteins, the selected genes had different putative functions including structural function (e.g., *cfiA* gene involved in membrane protein catalysis for ATP synthesis, transport and motility) (33) as well as DNA packaging and lysis (e.g., DNA braking-rejoining protein, lysozyme, prophage tail assembly protein) (Table 2).

Interestingly, the majority (n=7/8) of these predictors were located in mobile genetic elements regions such as putative prophages and plasmid. The presence of unique accessory genomic biomarkers within prophage regions has been recently shown to contribute to enhancing the ability of tracing-back the origin of epidemic clones of *S.* Typhimurium and its monophasic variant on a geographical scale (34). Here, half of the animal source genomic markers identified in the assessment of the probability of correct self-attribution were prophage-related genes, suggesting that prophage elements might play a crucial role also in source-tracking studies. As observed by Zhang et al. (17), plasmid and prophage-related loci may constitute highly informative predictors of livestock sources of *S*. Typhimurium and therefore contribute to the optimization of source attribution models for surveillance or outbreak investigations.

The importance of accessory genes in host adaptation is not limited to *Salmonella*. Recent studies focused on the pangenome of *Campylobacter jejuni* showed that host-segregating genomic factors located in the accessory genomes constitute epidemiological markers for source attribution (12; 35).

### Source of the *Salmonella* spp. environmental strains

*Salmonella* strains are frequently detected in surface water samples, e.g. 30.1% of samples from three French coastal catchments (31), 43% of samples from Georgia (USA) (36) and 23% of samples from Canada (37). Among *Salmonella* serovars, *Salmonella* Typhimurium and its monophasic variants were isolated more frequently in some sites than others (31). Pigs, poultry, and ruminants (e.g., cattle) constitute a relevant source of these pathogens acting as natural *Salmonella* reservoirs and without showing any symptoms while shedding into the environment (38). In this study, pig source was found to be the most likely source of contamination of most of the *S*. Typhimurium and *S*. 1,4,[5],12:i:-strains isolated in the environment, confirming the results from several sources tracking studies that pointed out some associations between environmental *Salmonella* spp. strains and pig production (39; 37). This is also consistent with the fact that pigs are asymptomatic carriers of *S*. Typhimurium generating a major reservoir for *Salmonella* contaminating the environment around the primary production sites (40; 41).

Although waterborne outbreaks of *Salmonella* have been rarely reported, a secondary waterborne outbreak has been linked to *S*. 1,4,[5],12:i:- and the leaching of animal faecal matter into groundwater destined for human consumption in a rural community (42). In addition, anthropological activities such as fertilization with animal manure and irrigation with water contaminated by livestock and/or wildlife faecal matter seem to be likely involved in *Salmonella* outbreaks and consumption of fresh produce (43; 44). Gaining insights into the animal sources of strains contaminating natural environments is therefore of great importance for providing evidence to support targeted interventions and policy development to reduce the public health risk.

## Author statements

### Author contributions

LG, MG, NM, TH, and FP conceived the study. SCS contributed to the selection of the data. SL and MLV performed the pre-sequencing steps. LG conceptualized algorithms. LG and FP implemented scripts and drafted the manuscript. All authors commented and approved the final manuscript, take public responsibility for appropriate portions of the content and agree to be accountable for all aspects of the work in terms of accuracy or integrity.

### Ethical approval

This study only included previously sequenced and published data. No new samples were collected for this study.

## Tables

Table 2: Predictors of animal sources. Number of isolates harboring the predictors with relative percentage of isolates from the different animal sources and environment and gene annotation from nucleotide and amino acid sequences obtain with Prokka and KEGG, respectively.

